# Phylogenetic evidence for independent origins of GDF1 and GDF3 genes in amphibians and mammals

**DOI:** 10.1101/293522

**Authors:** Juan C. Opazo, Kattina Zavala

## Abstract

Growth differentiation factors 1 (GDF1) and 3 (GDF3) are members of the transforming growth factor superfamily (TGF-β) that is involved in fundamental early-developmental processes that are conserved across vertebrates. The evolutionary history of these genes is still under debate due to ambiguous definitions of homologous relationships among vertebrates. Thus, the goal of this study was to unravel the evolution of the GDF1 and GDF3 genes of vertebrates, emphasizing the understanding of homologous relationships and their evolutionary origin. Surprisingly, our results revealed that the GDF1 and GDF3 genes found in amphibians and mammals are the products of independent duplication events of an ancestral gene in the ancestor of each of these lineages. The main implication of this result is that the GDF1 and GDF3 genes of amphibians and mammals are not 1:1 orthologs. In other words, genes that participate in fundamental processes during early development have been reinvented two independent times during the evolutionary history of tetrapods.

## Introduction

Growth differentiation factors 1 (GDF1) and 3 (GDF3) are members of the transforming growth factor superfamily (TGF-β) that were originally isolated from mouse embryonic libraries ^1,2^. They perform fundamental roles during early development, GDF1 has been mainly associated to the regulation of the left-right patterning, whereas GDF3 is mainly involved in the formation of the anterior visceral endoderm, mesoderm and the establishment of anterior-posterior identity of the body ^3–20^. Deficiencies in GDF1/GDF3 give rise to a broad spectrum of defects including right pulmonary isomerism, visceral *situs inversus*, transposition of the great arteries, and cardiac anomalies among others ^17,21–25^. In addition to their developmental roles, GDF1 has been described as a tumor suppressor gene in gastric cells, counteracting tumorogenesis by stimulating the SMAD signaling pathway ^26^, and GDF3 has been associated with the regulation of adipose tissue homeostasis and energy balance during nutrient overload ^27–29^.

The evolutionary history of the GDF1 and GDF3 genes is still a matter of debate due to the unclear definition of homologous relationships. Understanding homology is a fundamental aspect of biology as it allows us to comprehend the degree of relatedness between genes that are associated to a given phenotype in a group of organisms. This is particularly important for GDF1 and GDF3 as these genes perform biological functions during early stages of development that define key aspects of the body plan of all vertebrates ^3,4,13–20,5–12^. Until now, most of the inferences of the homology relationships of these genes have been based on functional information. For example, based on the developmental processes that these genes regulate, it has been suggested that the mammalian GDF1 gene is the true ortholog of the Vg1 (GDF1) gene found in amphibians ^15,17,21^. Furthermore, phylogenetic analyses performed by Andersson et al., (2007) were not able to define orthologous relationships between the GDF1 and GDF3 genes among vertebrates. However after performing genomic comparisons, these authors did propose that the GDF1 gene present in mammals is the true ortholog of the Vg1 (GDF1) gene present in amphibians ^30^. It was also suggested that the GDF3 gene could be an evolutionary innovation of mammals, as Andersson et al., (2007) did not find GDF3 sequences in the amphibian and bird genomes ^30,31^. Given this scenario, the single copy gene found in amphibians and birds would be a co-ortholog of the mammalian genes ^30^. More recently, a second copy of a GDF gene (derrière) has been annotated at the 3´ side of the Vg1 (GDF1) in the genome of the western clawed frog (*Xenopus tropicalis*), thus further complicating the definition of homologous relationships and the origin of genes that accomplish fundamental roles during early development in vertebrates.

With emphasis on understanding homologous relationships and their evolutionary origin, the goal of this study was to unravel the evolution of the GDF1 and GDF3 genes of vertebrates. Surprisingly, our phylogenetic analyses revealed that the GDF1 and GDF3 genes of amphibians and mammals are the products of independent duplication events of an ancestral gene in the ancestor of each of these groups. We also found the signature of two chromosomal translocations, the first occurred in the ancestor of tetrapods whereas the second was found in the ancestor of mammals. Thus, our results support the hypothesis that in amphibians and mammals, descendent copies of the same ancestral gene (GDF1/3) have independently subfunctionalized to perform key developmental functions in vertebrates.

## Results and Discussion

### Phylogenetic analyses suggest an independent origin of the GDF1 and GDF3 genes in mammals and amphibians

From an evolutionary perspective, the definition of homologous relationships among GDF1 and GDF3 genes is still a matter of debate ^30,31^. The resolution of this homology is important as these genes are involved in fundamental developmental processes which are conserved all across vertebrates ^8,32–34^. Thus, if extant species inherited these genes from the vertebrate ancestor, the developmental processes in which GDF1 and GDF3 are involved have a single evolutionary origin and are comparable among species.

Based on evolutionary analyses and the developmental processes in which GDF1 has been shown to participate, it has been suggested that GDF1 in mammals, birds, and amphibians are 1:1 orthologs ^15,17,21,30,35^. Given that it has not been possible to identify copies of the GDF3 gene in amphibians and birds, it has been proposed that this gene is an evolutionary innovation of mammals, and the single gene copy found in amphibians and birds is co-ortholog to the mammalian duplicates ^30^. However, a second copy of a GDF gene has been annotated in the genome of the western clawed frog (*Xenopus tropicalis*) ^36^. This newly described gene is located at the 3´ side of the GDF1 gene and indicates that amphibians, like mammals, also have a repertoire of two GDF genes (GDF1 (Vg1) and GDF3 (derrière)). This new finding further complicates the resolution of homologous relationships and the origin of genes involved in the early development of all vertebrates ^8,32–34^.

Our maximum likelihood and Bayesian reconstructions revealed that the GDF1 and GDF3 genes of amphibians and mammals were both reciprocally monophyletic (Fig. 1), indicating that these genes originated independently in each of these two lineages (Fig. 1). In agreement with the literature, we found only one gene in sauropsids (e.g. birds, turtles, lizards and allies), suggesting that they retained the ancestral condition of a single gene copy as found in non-tetrapod vertebrates (e.g. coelacanths, bony fish, chondricthyes and cyclostomes)(Fig. 1). Dot-plot comparisons provided further support for the presence of a single gene copy in sauropsids as no traces of an extra GDF gene were present in the syntenic region of representative species of the group (Fig. 2). However, we cannot rule out an alternative scenario of duplication and subsequent gene loss in the ancestor of sauropsids. Thus, our evolutionary analyses suggest that the GDF1 and GDF3 genes present in amphibians and mammals diversified independently in the ancestor of each of these lineages. In other words, genes that participate in fundamental processes during early development have been reinvented two independent times during the evolutionary history of tetrapods. As a consequence of the independent origin of these genes in amphibians and mammals, they are not 1:1 orthologs.

**Figure 1.**
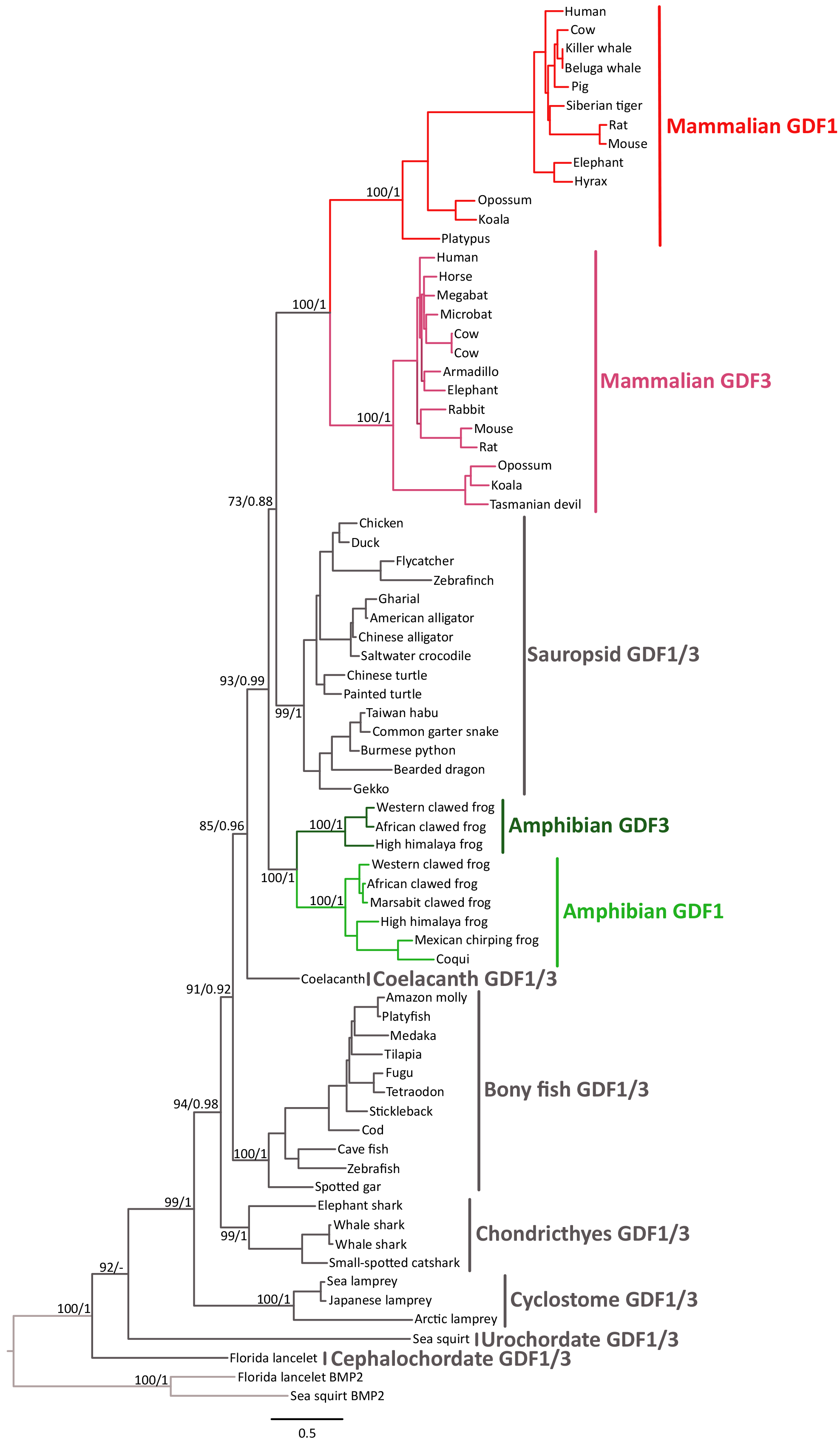
Maximum likelihood tree depicting evolutionary relationships among the GDF1 and GDF3 genes of chordates. Numbers on the nodes represent maximum likelihood ultrafast bootstrap and Bayesian posterior probability support values. BMP2 sequences from urochordates (*Ciona intestinalis*) and cephalochordates (*Branchiostoma floridae*) were used as outgroups.

**Figure 2.**
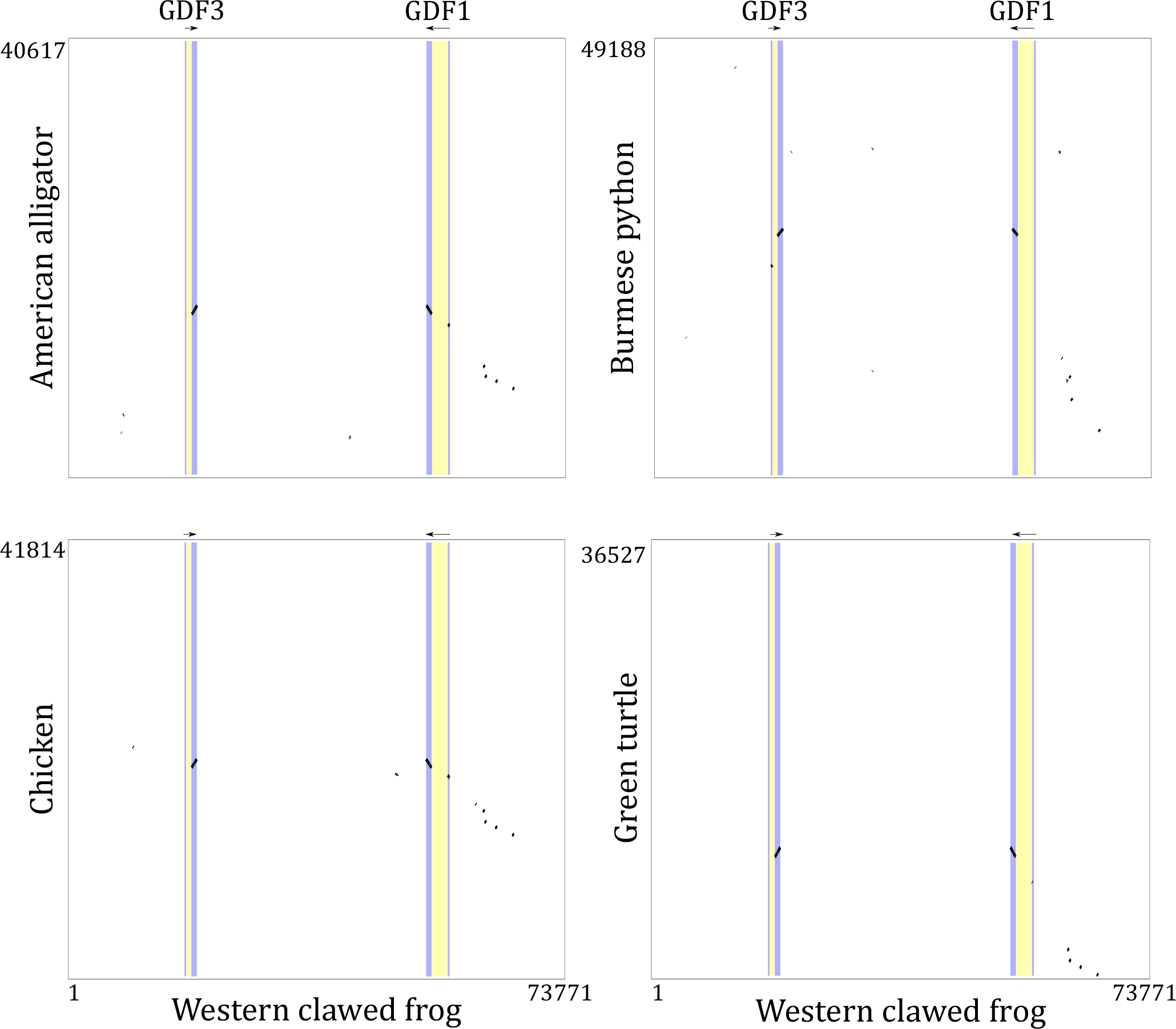
Dot-plots of pairwise sequence similarity between the GDF1 and GDF3 genes of the western clawed frog (*Xenopus tropicalis*) and the corresponding syntenic region in the American alligator (*Alligator mississippiensis*), Burmese python (*Python bivittatus*), chicken (*Gallus gallus*) and green turtle (*Chelonia mydas*).

To further test our hypothesis of the independent origins of the GDF1 and GDF3 genes in amphibians and mammals we performed topology tests. In these analyses we compared our phylogenetic tree (Fig. 1 and Fig. 3B) to the topology predicted from a one-duplication model in which the duplication event that gave rise to the GDF1 and GDF3 genes in amphibians and mammals occurred in the ancestor of tetrapods (Fig. 3A). In the one-duplication model, it is assumed that the ancestor of sauropsids had two gene copies and that subsequently one of these was lost. Thus, according to the one-duplication model (Fig. 3A), the predicted phylogeny would recover a monophyletic group containing GDF1 sequences from mammals, sauropsids, and amphibians sister to a clade containing GDF3 sequences from the same groups (Fig. 3A). Alternatively, the predicted phylogeny from the two-duplication model would retrieve a clade containing GDF1 and GDF3 sequences from mammals sister to a clade containing GDF1/3 sequences from sauropsids; additionally a clade containing GDF1 and GDF3 sequences from amphibians would be recovered sister to the mammalian/sauropsid clade (Fig. 3B). Results of the topology tests rejected the phylogeny predicted by the one-duplication model (Weighted Shimodaira and Hasegawa, p< 10^−4^; Weighted Kishino Hasegawa, p< 10^−4^). Thus, this result provided additional support to our hypothesis that the GDF1 and GDF3 genes of amphibians and mammals are the product of lineage independent duplication events in the ancestors of each of these groups.

**Figure 3.**
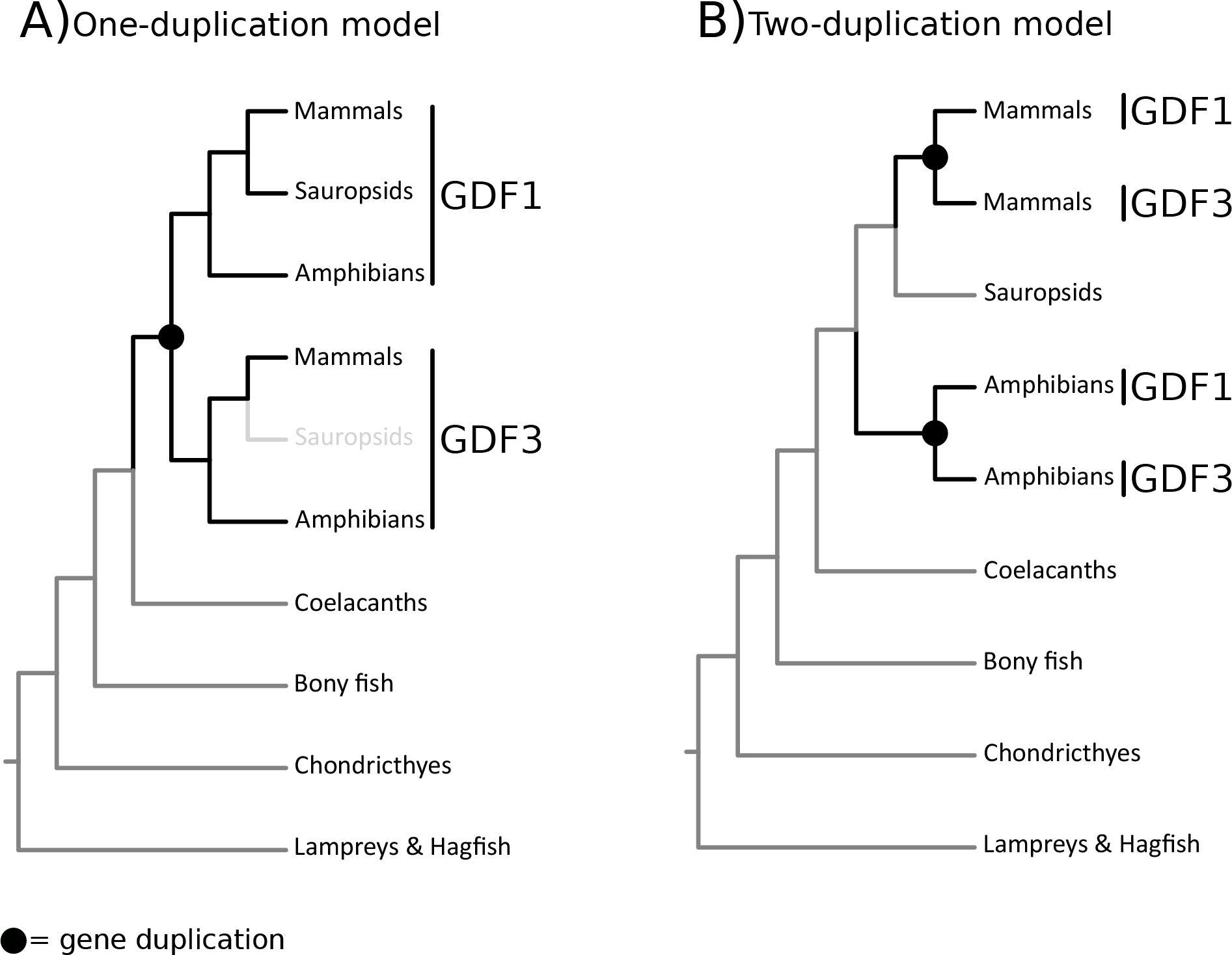
Schematic representations of alternative hypotheses of the sister group relationships among duplicated GDF genes in tetrapods. A) According to the one-duplication model the predicted phylogeny recovers a monophyletic group containing GDF1 sequences from mammals, sauropsids and amphibians sister to a clade containing GDF3 sequences from the same groups. B) The phylogenetic prediction from the two-duplication model retrieves a clade containing GDF1 and GDF3 sequences from mammals sister to a clade containing GDF1/3 sequences from sauropsids; additionally a clade containing GDF1 and GDF3 sequences from amphibians is recovered sister to the mammalian/sauropsid clade.

In the literature there are other cases of groups of genes that perform similar biological functions in a diversity of species but that have originated via lineage independent duplication events ^37–43^. Among these, the independent origin of the β-globin gene cluster in all main groups of tetrapods (e.g. therian mammals, monotremes, birds, crocodiles, turtles, squamates, amphibians) represents a well-documented phenomenon ^41–43^. This case is of particular interest as the two β-globin subunits that come from a gene family that has been reinvented several times during the evolutionary history of tetrapods are assembled in a tetramer with two α-globin subunits that belong to a group of genes that possess a single origin ^37,41–43^. Besides this example, the independent origin of gene families makes the task of comparison difficult, as the repertoire of genes linked to physiological processes in different lineages do not have the same evolutionary origin. Additionally, the fact that the evolutionary process can give rise to similar phenotypes following different mutational pathways makes the problem of comparing even more difficult (Natarajan et al., 2016). This is particularly important when extrapolating the results of physiological studies performed in model species to other organisms.

### Two translocation events during the evolutionary history of the GDF1 and GDF3 genes of tetrapods

Based on the chromosomal distribution of the GDF1 and GDF3 genes and the conservation pattern of flanking genes, we propose that during the evolutionary history of tetrapods, these genes underwent two chromosomal translocation events (Fig. 4). According to our analyses the genomic region that harbors the single gene copy of chondricthyes, bony fish, and coelacanths is conserved, yet this region in these groups differs from the regions in which GDF1 and GDF3 are located in tetrapods. Overall this suggests that the first translocation event occurred in the ancestor of tetrapods (Fig. 4). Interestingly, the genomic region where the single gene copy is located in non-tetrapod vertebrates, which is defined by the presence of upstream (BMP2, HAO1, TMX4 and PLCB1) and downstream genes (FERMT1, LRRN4, CRLS1), is conserved in tetrapods. Given this, it is possible to identify the chromosomal location where the gene was located in the tetrapod ancestor before the first translocation event (Fig. 4). On the other hand, the fact that the mammalian GDF1 and GDF3 genes are located in different chromosomes suggests that the second translocation event occurred in the ancestor of the group (Fig. 4). In humans, the GDF1 gene is located on chromosome 19 while GDF3 is located on chromosome 12. In opossum (*Didelphis virginiana*) the GDF1 and GDF3 genes are located on chromosomes 3 and 8, respectively.

**Figure 4.**
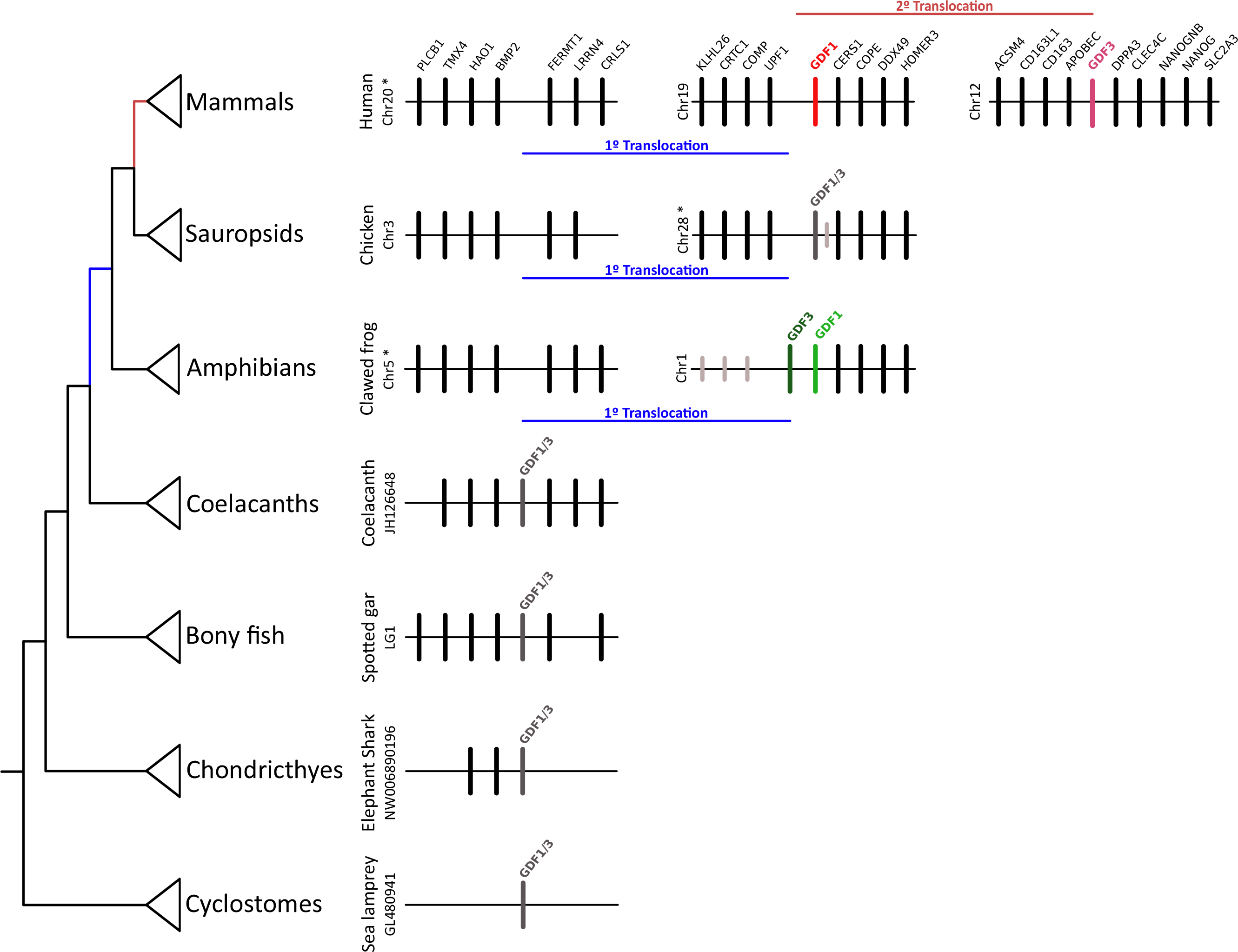
Structure of the chromosomal region containing the GDF1 and GDF3 genes of vertebrates. Asterisks denote that the orientation of the genomic piece is from 3’ to 5’, gray lines represent intervening genes that do not contribute to conserved synteny.

### Evolution of GDF1 and GDF3 genes in vertebrates

In this study we present compelling evidence suggesting that genes involved in the formation of the primitive streak, anterior visceral endoderm, mesoderm and the establishment of the left-right identity ^3,4,14–17,20,30,5–7,9–13^ in amphibians and mammals are the product of independent duplications events. As such, these results indicate that the GDF1 and GDF3 genes of amphibians and mammals are not 1:1 orthologs.

Thus, according to our results the last common ancestor of vertebrates had a repertoire of one gene (GDF1/3), and this condition is maintained in actual species of cylostomes, chondricthyes, bony fish, and coelacanths (Fig. 5). In the ancestor of tetrapods, the single gene copy was translocated from a chromosomal region defined by the presence of the BMP2, HAO1, TMX4, PLCB1, FERMT1, LRRN4, CRLS1 genes to a chromosomal region defined by the presence of the CERS1, COPE, DDX49 and HOMER3 genes (Fig. 4). After this translocation, the ancestral gene underwent a duplication event in the amphibian ancestor, giving rise to the GDF1 (Vg1) and GDF3 (derrière) genes as they are found in actual species. In the case of the western clawed frog (*Xenopus tropicalis*) these genes are located in tandem on chromosome 28 (Fig. 4). Sauropsids, the group that includes birds, crocodiles, turtles, lizards and snakes, inherited the ancestral condition of a single gene copy as is seen in non-tetrapod vertebrates (Fig. 5). Finally, in the ancestor of mammals, the ancestral gene also underwent a duplication event, giving rise to the GDF1 and GDF3 genes found in extant species of mammals. After duplication but before the radiation of the group, a second translocation event occurred in the ancestor of the group (Fig. 5). Thus, all mammals inherited a repertoire of two genes (GDF1 and GDF3) that are located on two chromosomes.

**Figure 5.**
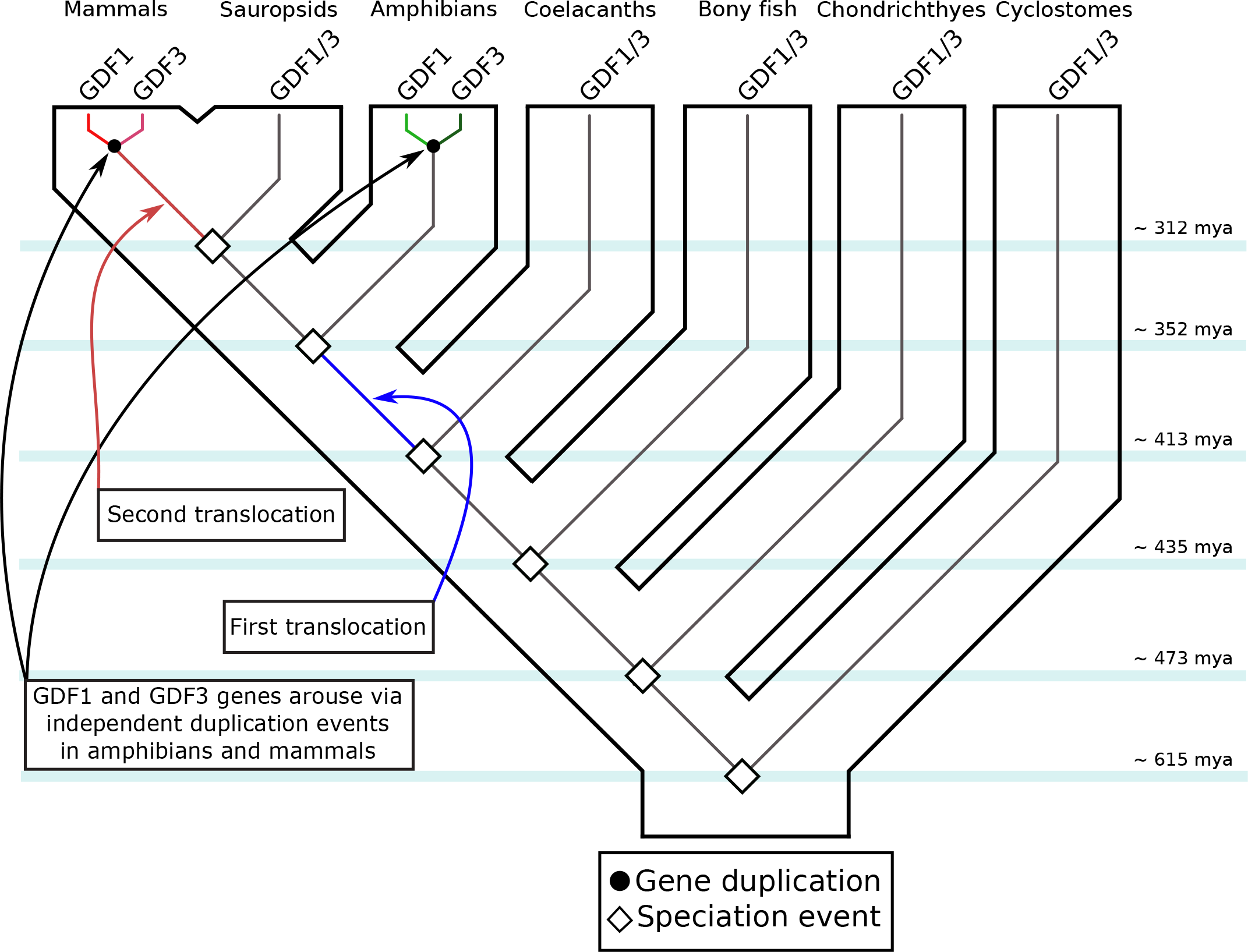
An evolutionary hypothesis of the evolution of the GDF1 and GDF3 genes in vertebrates. According to this model the last common ancestor of vertebrates had a repertoire of one gene (GDF1/3), a condition that has been maintained in actual species of cylostomes, chondrichthyes, bony fish, coelacanths and sauropsids. In the ancestor of tetrapods, the single gene copy was translocated to a different chromosomal location. After that, in the amphibian ancestor, the single gene copy underwent a duplication event giving rise to the amphibian GDF1 (Vg1) and GDF3 (derrière) genes. In the ancestor of mammals, the single gene copy also underwent a duplication event, giving rise to the mammalian GDF1 and GDF3 genes. After the duplication, but before the radiation of the group, a second translocation event occurred in the ancestor of the group. Thus, all mammals inherited a repertoire of two genes (GDF1 and GDF3) that were located on two chromosomes.

## Concluding remarks

This study provides a comprehensive evolutionary analysis of the GDF1 and GDF3 genes in representative species of all main groups of vertebrates. The main focus of this study was to unravel the duplicative history of the GDF1 and GDF3 genes and to understand homologous relationships among vertebrates. Understanding homology in this case is particularly important as these genes perform fundamental roles during early development that are conserved across vertebrates ^8,32–34^. Surprisingly, our results revealed that the GDF1 and GDF3 genes present in amphibians and mammals are the product of independent duplication events in the ancestor of each of these groups. Subsequently, the GDF1 and GDF3 genes of amphibians and mammals are not 1:1 orthologs. Our results also show that all other vertebrate groups - i.e non-tetrapods and sauropsids - maintained the ancestral condition of a single gene copy (GDF1/3).

From an evolutionary perspective the independent duplication events that occurred in the ancestors of mammals and amphibians could have resulted in the division of labor, with some degree of redundancy, of the function performed by the ancestral gene. In support of this idea, it has been shown that the GDF1 and GDF3 genes of mammals have partially redundant functions during development where GDF1 can to some degree compensate for the lack of GDF3 ^30^. Additionally, it has also been shown that double knockout animals (GDF1^−/−^ and GDF3^−/−^) present more severe phenotypes than those of either single knockout (GDF1^−/−^ or GDF3^−/−^)^30^. Detailed comparisons of the developmental roles of the GDF1 and GDF3 genes between mammals and amphibians will shed light regarding the reinvention of genes that possess fundamental roles during early development. On the other hand, comparing mammals and amphibians with sauropsids will provide useful information regarding the evolutionary fate of duplicated genes. Finally, it would be interesting to study the evolutionary history of genes that cooperate with GDF1 and GDF3 during early development (e.g. nodal) ^8,33,44^ in order to understand the evolutionary nature of the entire developmental network.

## Material and Methods

### DNA sequences and phylogenetic analyses

We annotated GDF1 and GDF3 genes in representative species of chordates. Our study included representative species from mammals, birds, reptiles, amphibians, coelacanths, holostean fish, teleost fish, cartilaginous fish, cyclostomes, urochordates and cephalochordates (Supplementary dataset 1 and 2). We identified genomic pieces containing GDF 1 and GDF3 genes in the Ensembl database using BLASTN with default settings or NCBI database (refseq_genomes, htgs, and wgs) using tbalstn (Altschul et al., 1990) with default settings. Conserved synteny was also used as a criterion to define the genomic region containing GDF1 and GDF3 genes. Once identified, genomic pieces were extracted including the 5´and 3´ flanking genes. After extraction, we curated the existing annotation by comparing known exon sequences to genomic pieces using the program Blast2seq with default parameters (Tatusova and Madden 1999). Putatively functional genes were characterized by an open intact reading frame with the canonical exon/intron structure typical of vertebrate GDF1 and GDF3 genes. Sequences derived from shorter records based on genomic DNA or cDNA were also included in order to attain a broad and balanced taxonomic coverage. Amino acid sequences were aligned using the L-INS-i strategy from MAFFT v.7 ^45^ (Supplementary Dataset 3). Phylogenetic relationships were estimated using maximum likelihood and Bayesian approaches. We used the proposed model tool from IQ-Tree ^46^ to select the best-fitting model (JTT+F+R5). Maximum likelihood analysis was also performed in IQ-Tree ^46^ to obtain the best tree. Node support was assessed with 1000 bootstrap pseudoreplicates using the ultrafast routine. Bayesian searches were conducted in MrBayes v.3.1.2 ^47^. Two independent runs of six simultaneous chains for 10×10^6^ generations were set, and every 2,500 generations were sampled using default priors. The run was considered to have reached convergence once the likelihood scores formed an asymptote and the average standard deviation of the split frequencies remained < 0.01. We discarded all trees that were sampled before convergence, and we evaluated support for the nodes and parameter estimates from a majority rule consensus of the last 2,000 trees. Sea squirt (*Ciona intestinalis*) and Florida lancelet (*Branchiostoma floridae*) BMP2 sequences were used as outgroups.

### Assessment of conserved synteny

We examined genes found up- and downstream of GDF1 and GDF3 in species representative of vertebrates. Synteny assessment were conducted for human (*Homo sapiens*), chicken (*Gallus gallus*), western-clawed frog (*Xenopus tropicalis*), coelacanth (*Latimeria chalumnae*), spotted gar (*Lepisosteus oculatus*), elephant shark (*Callorhinchus milii*) and sea lamprey (*Petromyzon marinus*). Initial ortholog predictions were derived from the EnsemblCompara database ^48^ and were visualized using the program Genomicus v91.01^49^. In other cases, the genome data viewer platform from the National Center for Biotechnology information was used.

## Acknowledgements

This work was supported by the Fondo Nacional de Desarrollo Científico y Tecnológico (FONDECYT 1160627) grant to JCO.

## Contributions

Designed the research: JCO; carried out the research: JCO and KZ; contributed materials/reagents/analysis tools: JCO; wrote the paper: JCO

